# Heat treatment functionalizes hepatocyte-like cells derived from human embryonic stem cells

**DOI:** 10.1101/2020.03.10.983130

**Authors:** Satoshi Imamura, Koki Yoshimoto, Shiho Terada, Ken-ichiro Kamei

## Abstract

Hepatocyte-like cells derived from human pluripotent stem cells (hPSC-HLCs) offer an alternative to primary hepatocytes commonly used for drug screenings and toxicological tests. Although tremendous efforts have been made to facilitate hepatic functions of hPSC-HLCs using growth factors and chemicals, these cells have not yet reached hepatic functions comparable to hepatocytes *in vivo*. Therefore, there exists a critical need to use an alternative trigger to facilitate hepatic functions in hPSC-HLCs. We noted that human liver temperature (around 39°C) is higher than normal human body temperature (around 36.5°C), yet hepatocytes are generally cultured at 37°C *in-vitro*. Here we showed that hepatic functions of hPSC-HLCs would be facilitated under physiological liver temperatures. We identified the optimal temperature by treating HLCs derived from H9 human embryonic stem cells (hESC-HLCs) at 39°C and 42°C. 42°C-treatment caused significantly greater cell death compared to 39°C. We also confirmed the increases of hepatic functions, such as secretion of albumin, cytochrome P450 3A4 (CYP3A4) activities, and collagen productions, without severe cell damages. To elucidate the underlying mechanisms of heat-induced hepatic functions, RNA-seq was to identify gene expression signatures due to 39°C-treated hESC-HLCs. This study also showed the possible mechanisms of heat-induced hepatic function via glucocorticoid receptor pathway and molecular chaperons. In combination with existing hepatic differentiation protocols, the method proposed here may further improve hepatic functions for hPSCs, and lead to the realization of drug discovery efforts and drug toxicological tests.

**Significance statement:** Hepatocyte-like cells derived from human pluripotent stem cells (hPSC-HLCs) offer an alternative to primary hepatocytes commonly used for drug screenings and toxicological tests. We noted that human liver temperature (around 39°C) is higher than normal human body temperature (around 36.5°C), affecting the in-vitro hepatic functions of hPSC-HLCs, such as metabolic activities. Here we showed that hepatic functions of hPSC-HLCs, albumin secretion, CYP3A4 activities, and collagen production would be facilitated under physiological liver temperatures at 39°C, without severe cell damages. RNA-seq was used to elucidate the underlying mechanisms of heat-induced hepatic functions. This study also showed the possible mechanisms of heat-induced hepatic function via glucocorticoid receptor pathway and molecular chaperons.

## Main text

### Introduction

Hepatocytes are the major cellular component of liver tissue and play an important role in protein synthesis/storage, carbohydrate metabolism, and production of cholesterol, bile acids, and phospholipids that contribute to homeostasis in vertebrates. The evaluation of drug candidates’ safety often focuses on vertebrate livers because hepatocytes metabolize chemical substances and drugs *in vivo* (1). Currently, primary human hepatocytes and cell lines (e.g., HepG2 and HepaRG) (2, 3) are used for liver research, but the former is difficult to obtain from healthy donors and maintain liver functions. The latter does not represent healthy liver function (e.g., drug metabolism and transport function) due to cancerous characteristics. Researchers have recently turned to stem cell technologies to respond to the need for alternative cost-efficient, homogenous, readily available, and viable *in-vitro* cells for liver research.

Hepatocyte-like cells (HLCs) derived from human pluripotent stem cells (hPSC-HLCs) have considerable potential to provide optimal hepatocyte function during drug screening and toxicity tests (4). Human pluripotent stem cells (hPSCs), embryonic stem cells (ESCs) (5), and induced pluripotent stem cells (iPSCs) (6) all can differentiate into almost any kind of cell and possess the ability to self-renew indefinitely. Previous studies have successfully induced the differentiation of hPSCs into hPSC-HLCs using growth factors and chemicals (e.g., hepatocyte growth factor (HGF), dexamethasone (DEX), oncostatin M (OSM)) (7–11), but these hepatocytes still have fetal liver functions. Especially, cytochrome p450 3A4 (CYP3A4), which is mainly produced in the mature liver and intestine and is known to be the most critical metabolic enzyme to consider when optimizing drug treatments, is low expression level but CYP3A7, which is the primary human fetal liver CYP3A, is still high expression level. Furthermore, albumin (ALB) secretion level has been improved, but alpha-fetoprotein (AFP), a fetal liver marker and is decreased in the adult liver, is still high. These studies demonstrate the need to develop new methods to obtain in-vitro hepatic function of hPSC-HLCs.

Previous research investigating methods to achieve the mature hepatic function of hPSC-HLCs *in-vitro* have not fully considered a human liver’s *in vivo* physiological conditions. By exploring the *in vivo* hepatic physiological conditions, we notice that the liver’s temperature is higher than the other body temperature or general *in-vitro* cell-culture conditions; this is because hepatocytes work as heat-producing organs during sugar, protein, and lipid metabolism (12, 13). On the other hand, since hPSC-HLCs do not reach hepatocytes’ functional levels *in vivo*, we hypothesize that hPSC-HLCs do not produce the necessary temperature for functioning. Indeed, other studies showed that higher temperatures (around 39°C) for *in-vitro* cell culture promote differentiation and growth of myoblasts and neurons in other heat-producing organs, such as muscle and brain, respectively (14, 15). In contrast, it has been suggested that spermatogenesis requires low temperature (16). Thus, each organ needs to be exposed with an appropriate temperature to maximize its function; there is a clear need to identify the proper temperature to functionalize hPSC-HLCs.

Here we demonstrate heat-induced functionalization of hPSC-HLCs and elucidate the functionalization process’s underlying mechanism by heat stimulation. We conclude that mild heat treatment at 39°C during the hepatocyte-maturation process, but not higher like 42°C, were able to functionalize hPSC-HLCs, such as albumin secretion, exogenous substances’ uptake, and excretion and metabolic activities, such as cytochrome P450 (CYP) family (e.g., CYP3A4 and CYP3A7), after 12-day treatment. We also show that induction of CYP3A7 is regulated by the expression of glucocorticoid receptor-induced by heat treatment. To further understand the molecular mechanisms, RNA sequencing (RNA-seq) and molecular biological analyses were conducted and reveal that heat treatment induced the critical genes associated with not only molecular chaperons of heat shock proteins (HSPs) but also extracellular matrices (ECMs) associated with liver structures and functions.

### Results

#### hESC-HLCs survive during 39°C-treatment

To prove that higher temperature than normal body temperature (37°C) functionalize hESC-HLCs, we treated the hepatic progenitor cells derived from WA09(H9) hESCs with higher temperatures at 37, 39, and 42°C during the hepatic differentiation process (17) from day 14 (**Fig. 1A**). 42°C-treated cells at 15 days after heat treatment (d.a.h.) showed apparent cell detachment from a cell-culture dish, while for 39°C-treated cells, there was no notable detachment and morphological change compared to 37°C-treated cells (**Fig. 1B**). Compared to 37°C-treatment (set as 100% cell survival), the numbers of 42°C-treated cells living at 6 and 15 d.a.h. were significantly reduced (54% ± 12% and 13% ± 12%, respectively) compared to that of 39°C-treated cells (82% ± 13% and 92% ± 8%, respectively; **Fig. 1C**). Therefore, we used only 37°C and 39°C-treatment to facilitate hepatic functions for hESC-HLCs.

**Figure 1.**
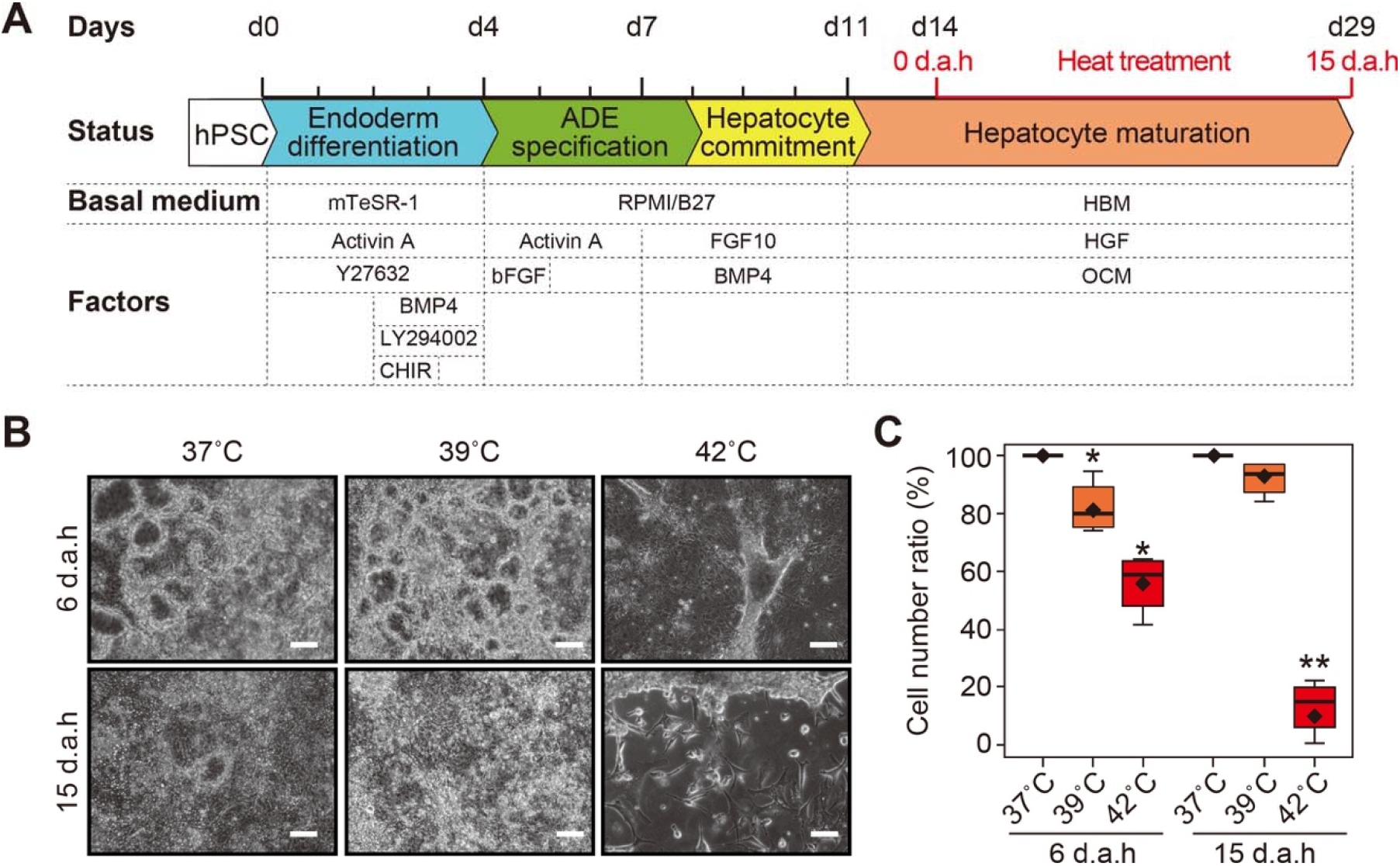
Functionalization of hepatocyte-like cells derived from hESCs by heat treatment. **A.** Schematic representation of the protocol to functionalize hESC-HLCs. DAH represents days after heat treatment. Initially, hPSCs were treated with mTeSR-1 medium supplemented with combinations of Activin A, BMP-4, CHIR99021 (CHIR), LY294002, and Y27632 to induce definitive endoderm (DE) differentiation until Day 4 (d4). Then, cells were treated with Activin A in RPMI medium supplemented with B27 for anterior definitive endoderm (ADE) specification until Day 8. As the next step, cells were treated with BMP-4 and FGF-10 for inducing hepatocyte commitment in RPMI medium supplemented with B27 until Day 11. Finally, cells were treated with oncostatin M (OSM) and hepatocyte growth factor (HGF) in basal hepatocyte medium for hepatocyte maturation. On day 14, cells were incubated at 37, 39, or 42°C, and half the total amount of the medium was changed every day. **B.** Microscopic images of hESC-HLCs treated with 37, 39, and 42°C at 6 and 15 d.a.h. Scale bars represent 100 μm. **c** Box plot showing the percentiles of living hESC-HLCs after heat treatments at 37, 39, and 42°C at 6 and 15 d.a.h. (n = 4). Centerlines of box plots indicate medians; box limits indicate the 25th and 75th percentiles as determined by *R* software; whiskers extend 1.5 times the interquartile range from the 25th and 75th percentiles; * = *P* < 0.05; each plot and error bar represents a mean ± SD.

#### 39°C-treatment facilitates hESC-HLCs’ hepatic functions

To evaluate the functionality of hESC-HLCs treated with 37°C and 39°C, we examined the effects on hepatic functions, such as albumin secretion, drug uptake/secretion, glucose storage, and molecular marker expression (**Fig. 2**).

**Figure 2.**
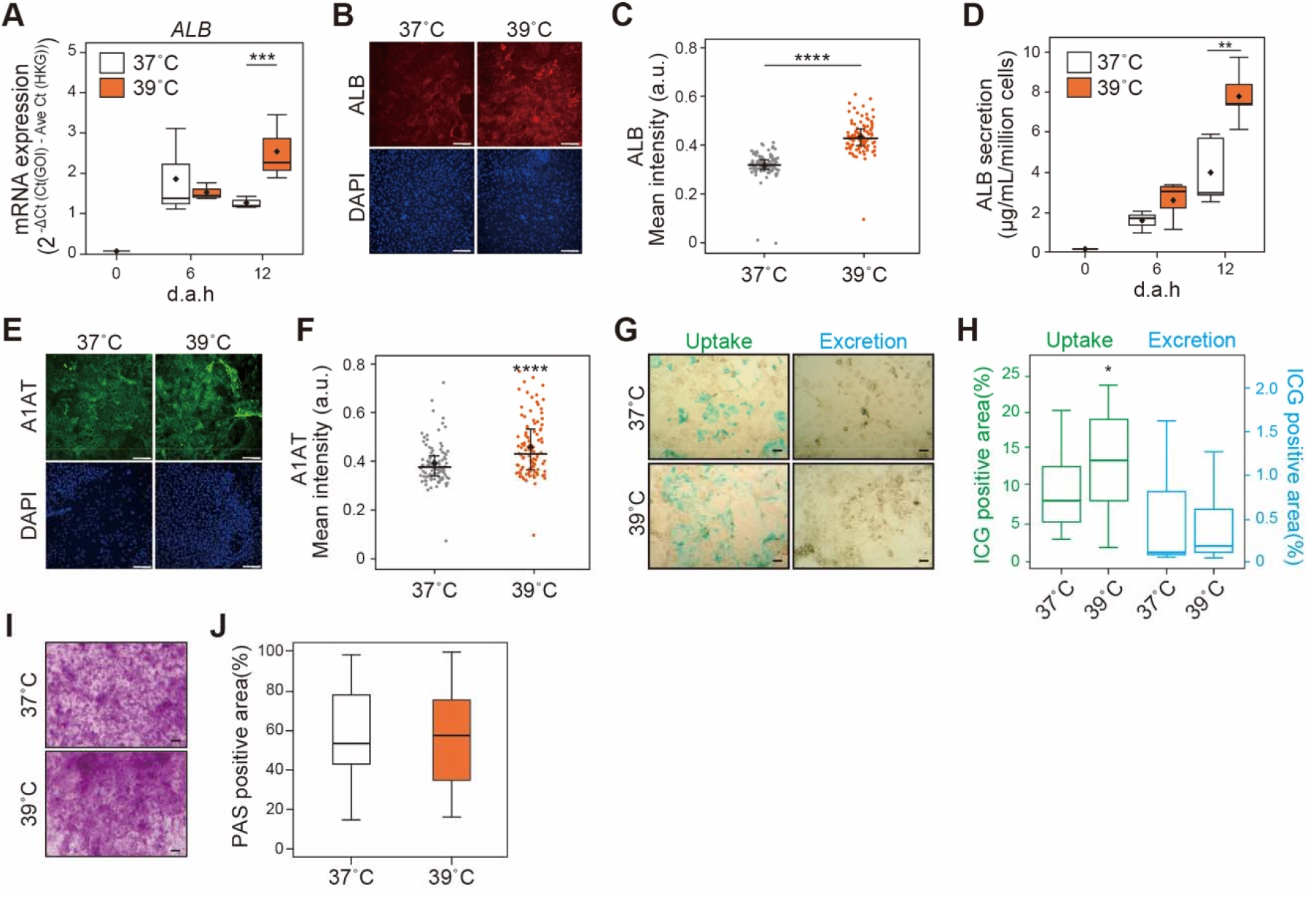
Heat treatment at 39°C facilitates hepatic functions of hESC-HLCs. (A) Gene expression of albumin (*ALB*) in 37°C- and 39°C-treated hESC-HLCs at 0, 6 and 12 d.a.h. evaluated by quantitative RT-PCR (n = 4). (B, C) Immunofluorescent micrographs (B) and single-cell profiling (C) of ALB in 37°C- and 39°C-treated hESC-HLCs at 12 d.a.h. DAPI was used for nuclei staining. Scale bars = 50 μm. (D) ALB secretion from 37°C- and 39°C-treated hESC-HLCs at 12 d.a.h. measured by ELISA. (E,F) Immunofluorescent micrographs (E) and single-cell profiling (F) of α1-anti trypsin (A1AT) in 37°C- and 39°C-treated hESC-HLCs at 12 d.a.h. DAPI was used for nuclei staining. Scale bars = 50 μm. (G, H) Micrographs (G) and the percentiles of positive area (H) of 37°C- and 39°C-treated hESC-HLCs at 12 d.a.h., stained with indocyanine green (ICG). Scale bars = 50 μm. (n = 20). (J, K) Micrographs (J) and the percentiles of positive area (K) of 37°C- and 39°C-treated hESC-HLCs at 12 d.a.h., stained with periodic acid schiff (PAS). Scale bars = 50 μm. (n = 20). In all panels, where applicable, center lines of box plots indicate medians; box limits indicate the 25th and 75th percentiles as determined by *R* software; whiskers extend 1.5 times the interquartile range from the 25th and 75th percentiles; \**p* < 0.05, \*\**p* < 0.01, \*\*\**p* < 0.005, \*\*\*\**p* < 0.0001; each plot and error bar represents a mean ± SD.

In comparison with 37°C-treated hESC-HLCs, 39°C-treated cells increased albumin (ALB) expression of both mRNA (p = 0.007, **Fig. 2A**) and protein (p < 0.001, **Fig. 2B** and **2C**). Importantly, 37°C- and 39°C-treated hESC-HLCs showed albumin secretion at 6 and 12 d.a.h., but at 12 d.a.h., 39°C-treated cells expression was significantly higher than those of 37°C-treated cells (p = 0.002, **Fig. 2D**). Similarly, the expression of α1anti-trypsin (A1AT, or called SERPINA1), one of liver maturation markers, was also increased in 39°C-treated cells (**Fig. 2E** and **2F**).

Indocyanine green (ICG) (18) was used for 37°C- and 39°C-treated hESC-HLCs to visualize the cells to confirm their abilities to uptake and excrete exogenous substances (**Fig. 2G** and **2H**). The ICG uptake levels of 39°C-treated cells were significantly higher than those of 37°C-treated cells at 12 d.a.h. (p = 0.039), but the excretion levels were not significantly different at 13 d.a.h. (p = 0.616). Since the transporters (e.g., solute carrier organic anion transporter family member 1B3 [*OATP1B3*] and Na^+^-taurocholate cotransporting polypeptide [*NCTP*, or known as sodium/bile acid cotransporter]) are involved in the uptake of ICG (19, 20), we confirmed their gene expression levels in both cells by quantitative RT-PCR. However, there was no significant difference neither (*SI Appendix*, **Fig. S1**).

To confirm the ability of glycogen storage of hESC-HLCs, periodic acid- Schiff (PAS) staining was performed (21) and indicated that both 37°C- and 39°C-treated hESC-HLCs could store glycogen in the cells (**Fig. 2I**). PAS-positive areas reached over 50% of these cells, but they did not show any significant difference (**Fig. 2J**).

#### 39°C-treatment activates hESC-HLCs’ CYP3A activity

To investigate the effects on the cytochrome P450 (CYP) family, which is responsible for metabolism, the expression of hepatic CYP genes (e.g., *CYP1A2, CYP2A6, CYP2C8, CYP2D6, CYP2E1, CYP3A4, CYP3A7*, and *CYP7A1*), were examined by quantitative RT-PCR (**Fig. 3A**). *CYP2A6, CYP2D6, CYP2E1, CYP3A4, CYP3A7*, and *CYP7A1* genes showed continuous increases hepatic differentiation process in 37°C- and 39°C-treated hESC-HLCs. Moreover, *CYP2A6, CYP3A7*, and *CYP7A1* genes were elevated in only 39°C-treated cells. Since the *CYP3A7* gene was increased by hepatic differentiation and 39°C-treatment, we conducted the bioluminescent-based CYP3A activity assay to confirm CYP3A activities during the hepatic differentiation process (**Fig. 3B**). Both 37°C- and 39°C-treated cells increased in CYP3A activities, and CYP3A activity in the 39°C-treated cells significantly increased at 6 d.a.h. (p = 0.045) and 12 d.a.h. (p < 0.001). The used CYP3A activity assay can detect both CYP3A4 and CYP3A7 and cannot distinguish them. Therefore, to elucidate which CYP3A4 or CYP3A7 showed the activities in 39°C-treated cells, fluorescent immunocytochemistry followed by quantitative single-cell profiling was performed (**Fig. 3C-3F**). Resultantly, 39°C-treated cells expressed both CYP3A4 and CYP3A7 proteins more than those of 37°C-treated cells (**Fig. 3D** and **3F**). Thus, it is likely both CYP3A4 and CYP3A7 showed metabolic activities. In CYP3A4, we could not see the difference in its gene expression between both temperatures; this means that the elevation of CYP3A4 proteins might involve the translational process, while the increase of CYP3A7 activities in 39°C-treated cells is regulated after the transcription step.

**Figure 3.**
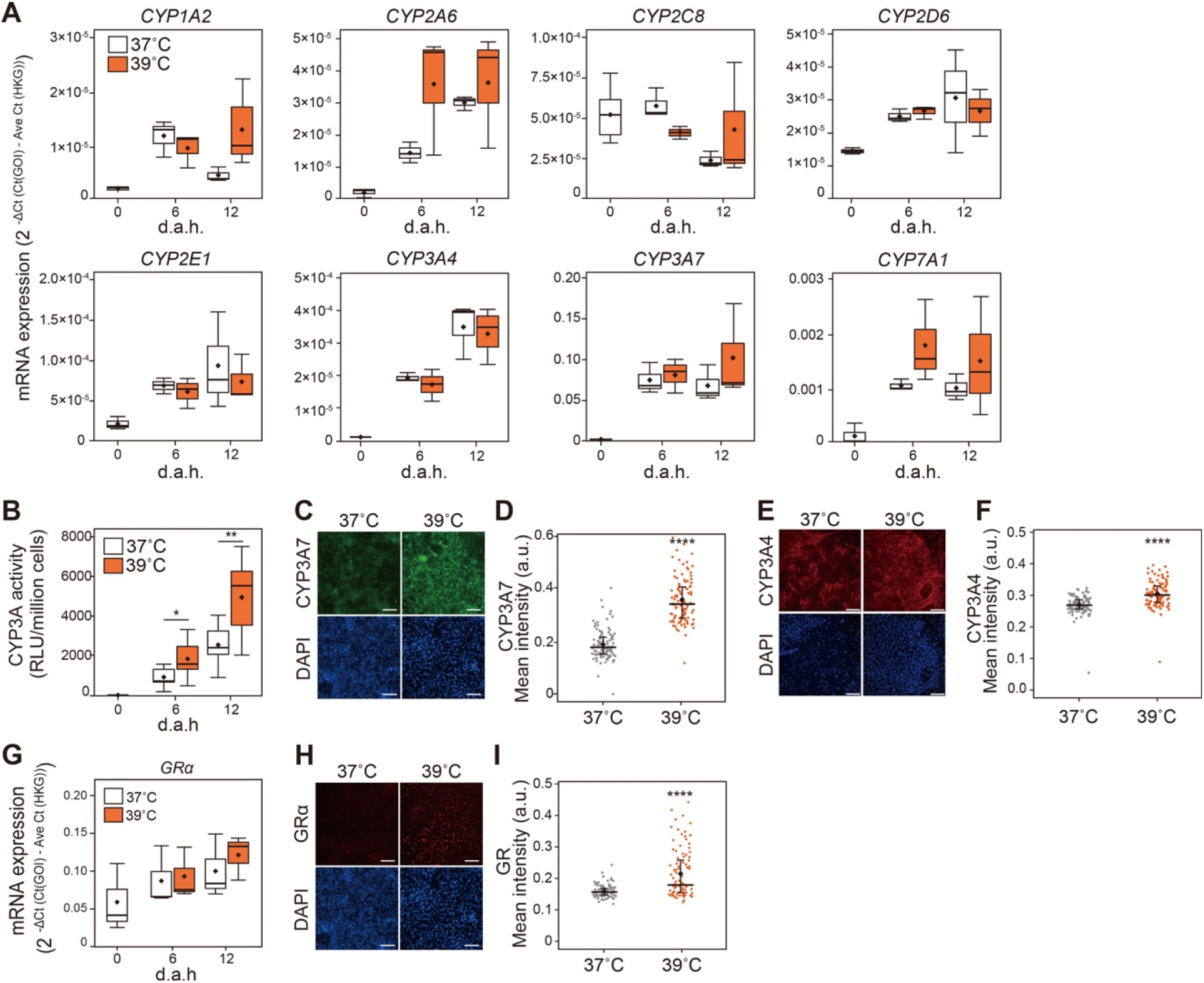
Evaluation of expression of hepatic metabolic CYP enzymes in hESC-HLCs with heat treatments. (A) Gene expression analysis of typical hepatic CYP enzymes, *CYP1A2, CYP2A6, CYP2C8, CYP2D6, CYP2E1, CYP3A4, CYP3A7* and *CYP7A1* during hepatic differentiation with heat treatments at 37°C and 39°C. (B) Bioluminescent-based CYP3A activity assay to confirm CYP3A activities during the hepatic differentiation process. (C, D) Immunofluorescent micrographs (C) and single-cell profiling (D) of CYP3A7 in 37°C- and 39°C-treated hESC-HLCs at 12 d.a.h. DAPI was used for nuclei staining. Scale bars = 50 μm. (E, F) Immunofluorescent micrographs (E) and single-cell profiling (F) of CYP3A4 in 37°C- and 39°C-treated hESC-HLCs at 12 d.a.h. DAPI was used for nuclei staining. Scale bars = 50 μm. (G) Gene expression analysis of glucocorticoid receptor (GR) during hepatic differentiation with heat treatments at 37°C and 39°C. (H, I) Immunofluorescent micrographs (H) and single-cell profiling (I) of CYP3A7 in 37°C- and 39°C-treated hESC-HLCs at 12 d.a.h. DAPI was used for nuclei staining. Scale bars = 50 μm. In all panels, where applicable, center lines of box plots indicate medians; box limits indicate the 25th and 75th percentiles as determined by *R* software; whiskers extend 1.5 times the interquartile range from the 25th and 75th percentiles; \**p* < 0.05, \*\**p* < 0.01, \*\*\**p* < 0.005, \*\*\*\**p* < 0.0001; each plot and error bar represents a mean ± SD.

To gain further insights into CYP3A7 regulations by heat treatment at 39°C, we searched the promoter region of CYP3A7 using GeneCards (22) and found that *CYP3A7* expression could be regulated by the glucocorticoid receptor (GR). It was also reported that GR played a central role in the transcriptional regulation of CYP3A7 (23). Therefore, we conducted quantitative RT-PCR analysis and fluorescent immunocytochemistry followed by quantitative single-cell profiling to evaluate GR expression in hESC-HLCs. *GR* mRNA expression was gradually increased during the hepatic differentiation process, but not significantly different between 37°C- and 39°C-treated cells (*p* = 0.521, **Fig. 3G**). On the other hand, GR protein was expressed in the nucleus of 39°C-treated hESC-HLCs, but not expressed in 39°C-treated cells (**Fig. 3H** and **3I**). These results indicate that the GR protein regulates the CYP3A7 transcription during heat treatment at 39°C.

#### Global transcriptional analysis identified gene expression signatures of 39°C-treated hESC-HLCs

To identify the genes affected by heat treatment at 39°C, RNA was harvested from hESC-HLCs at three time-points with three biological replicates (0, 6, and 12 d.a.h.; **Table S1**). We performed a time-course RNA-seq analysis using maSigPro (23) and identified 320 significant differentially expressed genes (DEGs, **Table S2**). Hierarchical clustering based on the identified genes was categorized into 5 clusters (**Fig. 4A, B** and *SI Appendix*, **Fig. S2-S3**). Cluster 1 has DEGs which the expression levels of 39°C-treated hESC-HLCs were similar to those of 37°C-treated cells until 6 d.a.h., and were significantly higher from 6 to 12 d.a.h. DEGs in Cluster 2 showed that both 37°C- and 39°C-treated hESC-HLCs increased the DEGs, but 39°C-treated hESC-HLCs have significantly higher expression during the hepatic differentiation process. On the other hand, Cluster 3 has DEGs, which continuously decreased their expression during hepatic differentiation, but does not show the difference between the temperatures. Cluster 4 is quite similar to Cluster 1. DEGs in Cluster 5 were unique since the DEGs showed increased from 0 to 6 d.a.h., once, but decreased from 6 to 12 d.a.h. Furthermore, 39°C-treated hESC-HLCs showed significantly higher expression of the DEGs in Cluster 5, compared with 37°C-treated cells for the tested period.

**Figure 4.**
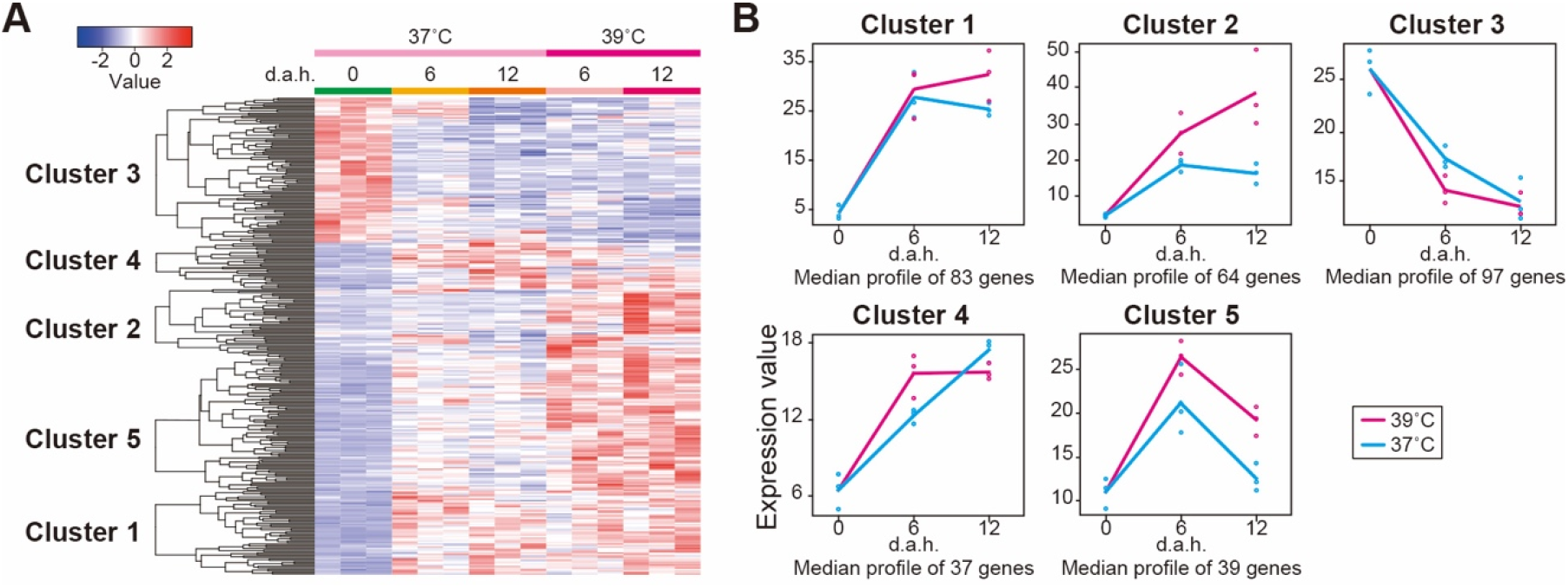
Global transcriptional analysis to identify the specific gene signatures in hESC-HLCs with heat treatments at 37°C and 39°C. (A) Hierarchical clustering and heat map of the differentially expressed genes in hESC-HLCs with heat treatments at 37°C and 39°C. (B) Typical plots of five clusters identified in (A). Each dot represents the median of expression values of each sample.

**Figure 5.**
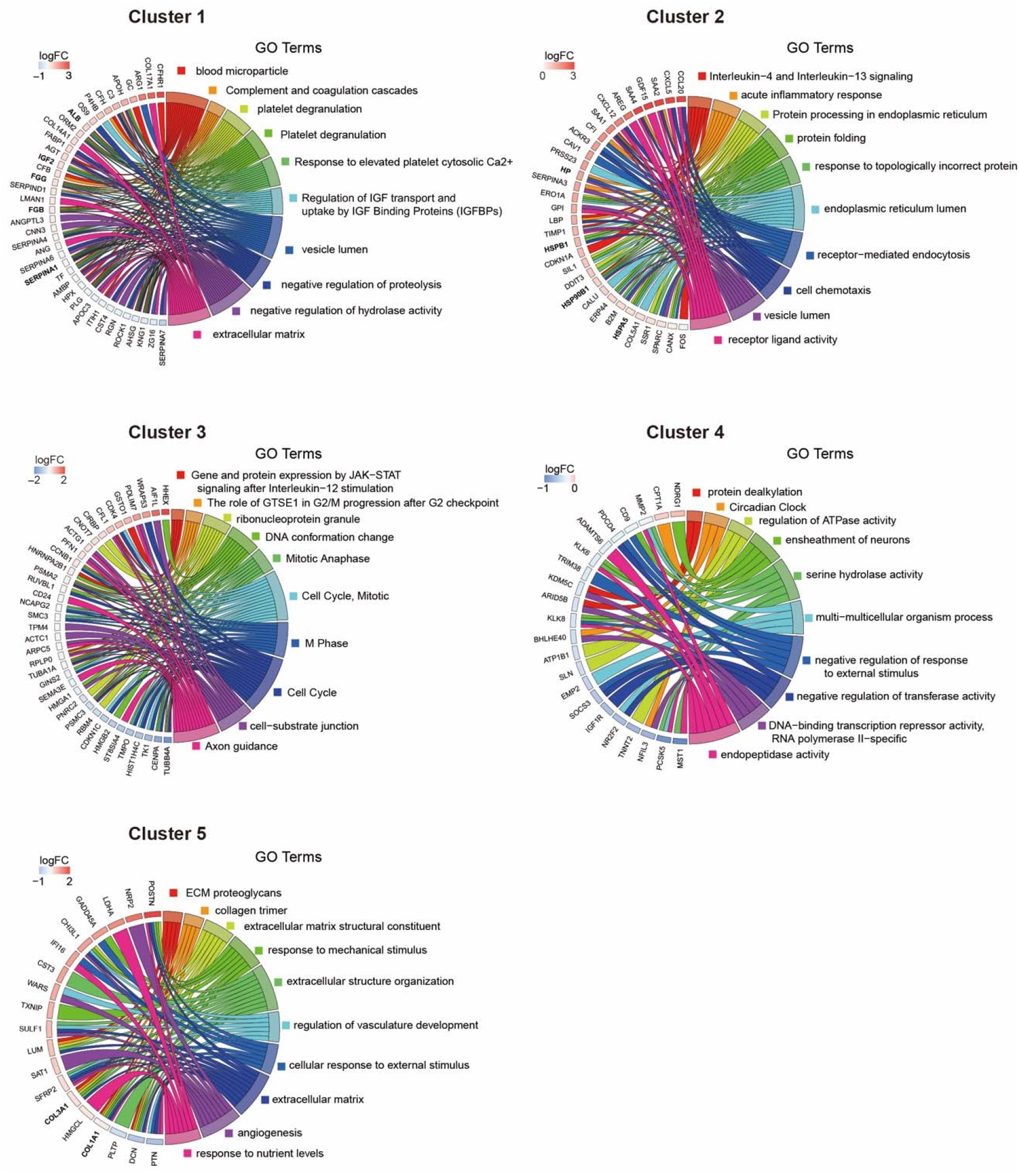
Chord diagram presenting enriched GO clusters for the differentially expressed genes hESC-HLCs with heat treatment at 37°C and 39°C. In each chord diagram, enriched GO clusters are shown (right), and genes contributing to this enrichment shown (left). Each cluster was found in Figure 4A.

To clarify what kind of go term or pathway the altered gene in each cluster, gene ontology (GO) enrichment, KEGG, and Reactome (24) analyses were performed (**Fig. 4**, *SI Appendix*, **Fig. S4**, and **Table S3**). In terms of Cluster 1, this has 83 genes, and particularly, complement factor H related 1 (*CFHR1*), *ALB*, insulin-like growth factor 2 (*IGF2*), fibrinogen beta chain (*FGB*), and fibrinogen gamma chain (*FGG*) had shown the high expression values. These genes showed biological terms associated with liver functions such as “blood microparticle” (GO:0072562), “Complement and coagulation cascades” (hsa04610), “platelet degranulation” (GO:0002576), and “Regulation of Insulin-like Growth Factor transport and uptake by Insulin-like Growth Factor Binding Proteins (IGFBPs)” (R-HSA-381426). They had a high enrichment ratio (FDR < 0.05). In light of the results shown in **Fig. 2**, We have validated the *ALB* gene expression and protein expression and secretion. Moreover, it was reported that CFHR1 was expressed specifically in an adult liver (25). Taken together, we found that the genes in Cluster 1 have roles of liver functions activated by heat treatment.

Cluster 2 has 64 genes, including haptoglobin (*HP*), heat shock protein 90kDa beta member 1 (*HSP90B1*), heat shock protein family A (HSP70) member 5 (*HSPA5*), and heat shock protein beta-1 (*HSPB1*). These genes were associated with the terms “acute inflammatory response” (GO:0002526), “Protein processing in endoplasmic reticulum” (hsa04141), and “protein folding” (GO:0006457) with high enrichment ratios (FDR < 0.05). Aforementioned, in this cluster, 39°C-treated hESC-HLCs facilitated the expression of the categorized genes for the tested period and might show the immune responses.

The terms and pathways in Cluster 3 were involved in cell division, such as “The role of GTSE1 in G2/M progression after G2 checkpoint” (R-HSA-8852276), “Mitotic Anaphase” (R-HSA-68882), “Cell Cycle, Mitotic” (R-HSA-69278), and “M Phase” (R-HSA-68886) (FDR < 0.05). Since it has been reported that hepatocytes’ mitotic function decreases with maturation (26), our results suggested that cells treated at 39°C reduced the mitotic functions toward improved hepatic maturity.

Cluster 4 has terms and pathways, such as “protein dealkylation” (GO:0008214) and “Circadian Clock” (R-HSA-400253) with a high enrichment ratio. However, since the FDR values were above 0.05 (**Fig. S3**), they had less possibility to be activated by heat treatments.

Cluster 5 has the 39 genes with high expression values in 39°C-treated hESC-HLCs than 37°C-treated hESC-HLCs, such as collagen type I alpha 1 chain (*COL1A1*), collagen type III alpha 1 chain (*COL3A1*). These genes are involved in ECM construction, such as “ECM proteoglycans” (R-HSA-3000178), “collagen trimer” (GO:0005581), and “extracellular matrix structural constituent” (GO:0005201) (FDR < 0.05).

#### 39°C-treatment activates collagen production of hESC-HLCs

Collagens are known to be a constitutive ECM of the liver and have various roles not only liver in development (27) but also in diseases (27–29). mRNA-seq results showed that heat treatment at 39°C increased the expression of *COL1A1* and *COL3A1*. Notably, ECM has been reported to promote hepatocytes’ functions such as CYP3A4, CYP3A7, and albumin (30, 31). Therefore, we performed fluorescence immunocytochemistry and quantitative single-cell profiling of COL1A and COL3A, and COL4A, which was not detected as DEGs by RNA-seq but is involved in the liver constitution. The expression of COL1A and COL4A were observed in both 37°C- and 39°C-treated hESC-HLCs, but 39°C-treated cells significantly higher than 37°C-treated cells (p < 0.001) (**Fig. 6A**–**6D**). On the other hand, COL3A had a low expression level in both 37°C- and 39°C-treated cells, and there was no significant difference (**Fig. 6E and 6F**). Thus, we confirmed that 39°C-treated hESC-HLCs elevated the expression of proteins of COL1A and COL4A.

**Figure 6.**
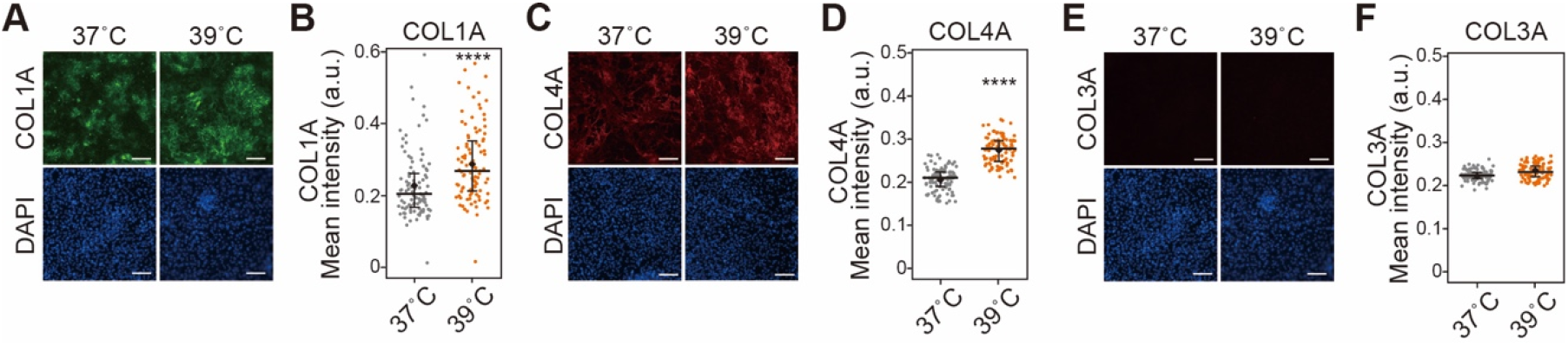
Evaluation of expression of collagen type I (COL1A), collagen type IV (COL4A), and collagen type III (COL3A) in hESC-HLCs with heat treatments. (A, B) Immunofluorescent micrographs (A) and single-cell profiling (B) of COL1A in 37°C- and 39°C-treated hESC-HLCs at 12 d.a.h. (C, D) Immunofluorescent micrographs (C) and single-cell profiling (D) of COL4A in 37°C- and 39°C-treated hESC-HLCs at 12 d.a.h. (E, F) Immunofluorescent micrographs (E) and single-cell profiling (F) of COL3A in 37°C- and 39°C-treated hESC-HLCs at 12 d.a.h. In all panels, where applicable, center lines of box plots indicate medians; box limits indicate the 25th and 75th percentiles as determined by *R* software; whiskers extend 1.5 times the interquartile range from the 25th and 75th percentiles; \**p* < 0.05, \*\**p* < 0.01, \*\*\**p* < 0.005, \*\*\*\**p* < 0.0001; each plot and error bar represents a mean ± SD. DAPI was used for nuclei staining. Scale bars = 50 μm.

### Discussion

In this study, we present the new way to functionalize HLCs derived from hPSCs by heat treatment at 39°C, but not a higher temperature like 42°C, and with chemical treatments. In general, while normal body temperature showed around 37°C, a core body temperature often showed ≥38.3°C, and the liver particularly showed such temperature because of its role as a heat-producing organ (14, 15). In contrast, such a high temperature from the range between 41 to 45°C is used as hyperthermia for cancer therapy because it often causes cell death for cancer cells and normal tissue cells (32, 33). We also frequently get a fever at temperatures over 39°C by various factors, causing extensive tissue damages (34). Thus, temperature control is a critical factor for proper functionality from cells to the whole body, and we need to investigate the optimal temperature for our target.

Regarding thermal effects on the survival of hESC-HLCs, 39°C-treatment showed a similar rate with 37°C conventional cell-culture setting, but 42°C- treatment reduced cell numbers. There are a couple of reasons for the reduction of cell numbers. The one is cell detachment. It has been reported that heat stress decreases integrin, a protein involved in cell adhesion, on the cell surface, causing cells to detach from the substrate (35). The other is heat-induced apoptosis. It has been reported that heat stress induces apoptosis through changes in mitochondrial membrane potential, activation of caspases, and activation of p53 (36). These reports and our results suggested that cell treatment at 42°C was not suitable for hepatic differentiation.

We found the elevation of ALB expression and A1AT and the uptake of exogenous substance, ICG (Fig. 2), by exploring the heat-induced hepatic functions. In the cases of ALB and A1AT, both of them were increased at both gene and protein levels. To gain deeper insights into their expression’s molecular basis, we explored the promoter region of their associated genes by using the GeneCards (37) and found that the *ALB* and *A1AT* promoters have the bonding domain TATA-binding protein (TBP). TBP was known as the target of heat shock protein 1 (HSF1) to work as transcription factors complex (38), and thus, the expression of ALB and A1AT might be regulated by HSF1-TBP complex heat treatment. It is important to note that HSF1 was reported to regulate GR’s expression (39, 40), and we observed that the gene and protein expressions of GR were increased by 39°C-treatment, leading to the induction of hepatic metabolic enzymes (CYP3A4 and CYP3A7). Although we could not see the difference in gene expression of HSF1 by heat treatment confirmed by RNA-seq, HSF1 would have an essential role in gene and protein regulation during 39°C-treatment. In terms of ICG uptake, primary transporters’ gene expression to uptake ICG into cells (e.g., *OATP1B3* and *NCTP*) did not show any difference by heat treatment. Therefore, we still need further investigation of the uptake mechanisms.

Interestingly, we found that 39°C-treated hESC-HLCs expressed COL1A1 and COL4A1 to generate a suitable environment for hepatic functions (**Fig. 6A-D**). ECMs have important roles for both liver development (27) and diseases (27–29), and each organ has unique ECM compositions and structures (41). In COL1A1, the promoter of COL1A1 has both GR and HSF1 binding domains found by GeneCards; GR and HSF1 might also regulate COL1A1’s gene expression during 39°C-treatment. In contrast, COL4A1 did not show an increase in its gene expression, but its protein expression showed a dramatic increase. That could be regulated by translation. Recently, it was reported that HSF1 had a role in ECM remodeling in cancer (42), and HSF1 could be involved in the increase of COL4A1 protein production.

Besides hepatocyte-associated genes, the genes of several molecular chaperones HSPs (Heat shock protein [HSP] 90 beta family member 1 [*HSP90B1*], HSP family A [HSP70] member 5 [*HSPA5*], and HSP family B [small] member 1 [*HSPB1*]) were induced by heat treatment at 39°C. Molecular chaperons generally work for protein folding, liver development (43, 44), and disease progression (45–47). Particularly, HSP90B1 and HSPA5 (or called glucose-regulated protein 78 [GRP78]) were expressed in the liver, according to the Human Protein Atlas (48–50). HSPB1 expression has been reported in a variety of organs, including the liver. In terms of cellular status, which we observed, 39°C-treated hESC-HLCs did not show cell apoptosis and cell growth progression similar to 37°C-treated hESC-HLCs. In contrast, we found that CYP3A4 activities were upregulated, although its gene expression did not change. These results suggested that molecular chaperons support CYP3A4 fording, resulting in the promotion of its activity. Thus, the upregulated chaperons worked for supporting protein folding rather than cancer- or disease-like status.

RNA-seq analyses also suggested the induction of genes associated with immune responses (e.g., serum amyloid A1 [*SAA1*], *SAA2*, and *SAA4*) by 39°C-treatment. SAA family is major acute-phase proteins produced during acute-phase response for inflammation and produced by hepatocytes or inflammatory cells, such as macrophages (51). Importantly, the SAA family was reported to induce anti-inflammatory M2-like macrophages during liver inflammation (52), and M2 macrophages have essential roles in liver repair and regeneration via cross-talks between M2 macrophages and hepatocytes (53, 54). In our study, macrophages were not incorporated in our cell-culture experiments, but it would be critical and interesting to investigate whether the incorporated macrophages show liver regeneration or cause inflammation.

Although we improved the functionality of hepatocytes derived from hPSCs, there is still a big room to improve more. Indeed, the obtained hepatocytes have not reached the adult liver stage since CYP3A7 and AFP, expressed during the fetal hepatic developmental process (55–57), were expressed in 39°C-treated hESC-HLCs. Therefore, additional treatments need to be established for the further maturation process of hepatocytes. Aforementioned, while chemical environmental cues have been studied to functionalized hPSC-HLCs, physical cues are still largely uncovered. To recapitulate liver-like cell aggregates, liver organoid technology with the use of hPSCs recently shows its strong potential for applications in drug discovery and regenerative medicine (58, 59). However, even such organoids, which are generally obtained with chemical environmental cues, do not reach the liver’s functional levels *in vivo*. Thus, physical environmental cues, such as shown in this study, would facilitate hepatocytes and liver organoids’ hepatic functions. Particularly, the thermal conditions for liver functionalization are quite interesting but not fully understood yet. We could not answer whether hepatocytes generate the heat first, or high environmental temperature allows hepatocytes to generate the heat, like a “chicken-and-egg” problem. In this study, we applied external heat treatment on hPSC-HLCs and confirmed that hPSC-HLCs were able to be functional, but we could not confirm whether hPSC-HLCs generated heat or not, since if so, that would be very small temperature changes. Recently, the intracellular thermal biosensors based on genetically engineered green fluorescent proteins have been established (60–62), the use of such biosensors would be beneficial to investigate the mechanisms of thermal regulation for liver functionalization.

Recently, organs-on-a-chip (OoC) or microphysiological systems (MPSs), based on microfluidic technology, show their great potential to recapitulate pathophysiological conditions *in-vitro* for applications in drug screening and toxicological testing and to reduce the use of experimental animals (63–67). Particularly, Liver-on-a-Chip technology is beneficial over the conventional cell-culture setting to apply three-dimensional (3D) well-designed cellular microenvironments and to promote hepatic functionalities *in-vitro* (68–70). In combination with this technology, we expect that our approach with heat treatments will facilitate the hepatic functions more. However, it will also raise a new concern. OoC platforms often have multiple tissue cells to investigate the inter-tissue interactions (67, 71), but each organ might require an optimal temperature to express its proper functions. Until now, to the best of our knowledge, any OoC or MPSs with multiple tissue cells are generally placed in the same incubator to carry out their functional assays, and there are no temperature controls for each organ. Microfluidic technology has great compatibility with micro-electrical mechanical systems (MEMS), and to solve the issues of temperature differences, MEMS-based thermal management systems (72, 73) should be integrated into OoC or MPSs for precise temperature controls for each organ.

### Conclusions

In conclusion, we showed the physiologically relevant heat condition at 39°C allows functionalizing hESC-HLCs, such as albumin secretion, CYP3A4 activities, and collagen productions, without severe cell damages. Since this approach is straightforward and does not require any special instruments, it would be rapidly used for practical applications of functionalized hPSC-HLCs for drug discovery and toxicological tests.

### Materials and methods

#### hESC culture

hESCs were used according to the guidelines provided by the ethical committee of Kyoto University. WA09(H9) (RRID:CVCL_9773, hPSCreg Name WAe009-A) hESCs used in this study were purchased from WiCell Research Institute (Madison, WI, USA). Before culturing, hESC-certified Matrigel (Corning, Corning, NY, USA) was diluted with Dulbecco’s modified Eagle medium (DMEM)/F12 medium (Sigma-Aldrich, St. Louis, MO, USA) at a 1:75 (v/v) ratio and coated onto a culture dish. The Matrigel was incubated in the culture dish for 24 h at 4°C. Then, excess Matrigel was removed, and the coated dish was washed with fresh DMEM/F12 medium. We used mTeSR-1-defined medium (Stem Cell Technologies, Vancouver, Canada) for daily culturing of hPSCs. For passaging, cells were dissociated with TryPLE Express (Thermo Fisher Scientific, Tokyo, Japan) for 3 min at 37°C and then harvested. A cell strainer was used to remove undesired cell aggregates from the cell suspension, and cells were then centrifuged at 200 × *g* for 3 min and resuspended in mTeSR-1 medium. Live/dead cells were counted using a NucleoCounter NC-200 (Chemetec, Baton Rouge, LA, USA). We used mTeSR-1 medium containing 10 μM of the ROCK inhibitor Y-27632 (Wako, Osaka, Japan) to prevent apoptosis of dissociated hPSCs on day 1. On subsequent days, we used mTeSR-1 medium without the ROCK inhibitor with daily medium changes.

#### Hepatic differentiation from hPSC

Before inducing differentiation, we coated a cell-culture dish with 0.1% gelatin in phosphate-buffered saline (PBS, Thermo Fisher Scientific) for 30 min at 25°C. We then aspirated the gelatin solution and introduced a DMEM/F12 medium (Sigma-Aldrich) onto the culture dish for serum coating at 37°C for 24 h. The medium was supplemented with 10% (v/v) fetal bovine serum (Cell Culture Bioscience, Tokyo, Japan), penicillin/streptomycin (Wako), and 100 μM β-mercaptoethanol (Sigma-Aldrich). The coated dish was then rinsed with fresh medium.

Cultured hPSCs were washed with PBS and treated with TryPLE Express at 37°C for 5 min, followed by the addition of basal medium and the transfer of the cell suspension into a 15 mL tube to induce endoderm differentiation. Cells were centrifuged at 200 × *g* for 3 min, the supernatant was removed, and then the cells were resuspended in mTeSR-1 medium supplemented with 10 μM Y27632 and 100 ng mL ^−1^ activin A (R&D Systems, Minneapolis, MN, USA), plated on a serum-coated culture dish, and cultured in a humidified incubator at 37°C with 5% CO_2_ for 24 h. At the end of day 1, the medium was replaced with fresh mTeSR-1 medium supplemented with 10 μM Y27632 and 100 ng mL^−1^ activin A and cultured for another 24 h. On day 2, the medium was replaced with mTeSR-1 medium supplemented with 10 μM Y27632, 100 ng mL^−1^ activin A, 10 ng mL^−1^ BMP-4 (R&D Systems), 10 μM LY294002 (Cayman Chemical, Ann Arbor, MI, USA), and 3 μM CHIR99021 (Stemgent, Cambridge, MA, USA), and cells were incubated for 24 h. On day 3, the medium was replaced with mTeSR-1 medium supplemented with 10 μM Y27632, 100 ng mL^−1^ activin A, 10 ng mL ^−1^ BMP-4, and 10 μM LY294002, and cells were incubated for 24 h. On day 4, the medium was replaced with Roswell Park Memorial Institute (RPMI) medium (Thermo Fisher Scientific) supplemented with B-27 (Thermo Fisher Scientific), 100 ng mL^−1^ activin A, and 100 ng mL^−1^ basic FGF, and cells were incubated for 24 h. To induce ADE specification, cells were treated with RPMI medium supplemented with 50 ng mL^−1^ activin A, with daily medium changes for three days. Cells were then treated with RPMI medium supplemented with 20 ng mL^−1^ BMP-4 and 10 ng mL^−1^ FGF-10 (R&D systems), with daily medium changes for four days. On day 12, the medium was replaced with hepatocyte-maturation medium (hepatocyte basal medium (Lonza, Basel, Switzerland) supplemented with 30 ng mL^−1^ oncostatin M (R&D Systems), 50 ng mL^−1^ HGF (PeproTech, Rocky Hill, NJ), and 25 mM HEPES (Wako) to induce maturation of the differentiated hepatocytes. On day 14, cells were incubated at 37, 39, or 42°C, and half the total amount of the medium was changed every day.

#### CYP3A activity assay

We used a cytochrome P450 3A (CYP3A) Assay and Screening System with Luciferin-IPA (Promega, Madison, MI, USA) to assess CYP3A activity. Samples were treated with a luciferin-IPA substrate (1:1000) in hepatocyte-maturation medium at 0, 6, and 12 d.a.h. of either 37°C or 39°C. We collected the medium after 1 h and added Luciferin Detection Reagent (Promega). After 15min, the CYP3A activity was measured in each sample with a Synergy HTX multi-mode reader (BioTek Instruments, Inc. Winooski, VT, USA). The total number of cells normalized the activity.

#### ELISA for human albumin

The medium cultured with cells were collected and stored at −80°C until use. The concentration of albumin secreted into the medium was measured using human albumin ELISA kit (Abcam, Cambridge, Cambridgeshire, UK, ab179887), following the manufacture’s protocol.

#### Immunocytochemistry

Cells were fixed with 4% paraformaldehyde (Wako, 161-20141) in PBS for 20 min at 25°C and then permeabilized with 0.1% Triton X-100 (MP Biomedicals, CA, USA) in PBS for 10 min at 25°C. Subsequently, cells were blocked in blocking buffer (5% normal goat serum, Vector; 5% normal donkey serum, Wako; 3% bovine serum albumin, Sigma-Aldrich; and 0.1% Tween-20, Nacalai Tesque, Inc, Kyoto, Japan) in PBS at 4°C for 16 h and then incubated at 4°C for 16 h with the primary antibody (anti-human CYP3A7 rabbit IgG, 1:500, Proteintech, Chicago, USA; anti-human A1AT rabbit IgG, 1:800, Dako, Tokyo, Japan. A0012; anti-human CYP3A4 mouse IgG, 1:25, Santa Cruz Biotechnology, Inc, CA, USA, sc-53850; and anti-human ALB mouse IgG, 1:50, R&D Systems, 188835) in blocking buffer. Cells were then incubated at 37°C for 60 min with a secondary antibody (AlexaFluor 488 Donkey anti-rabbit IgG, 1:1000, Jackson ImmunoResearch, PA, USA 711-546-152 and AlexaFluor 647 Donkey anti-mouse IgG, 1:1000, Jackson ImmunoResearch, 715-606-150) in blocking buffer before a final incubation with 4’,6-diamidino-2-phenylindole (DAPI; Wako 342-07431) at 25°C for 30 min.

#### Indocyanine green (ICG) uptake/excretion assay

Briefly, 1 mg mL^−1^ ICG (Sigma-Aldrich) was dissolved in the hepatocyte-maturation medium. Cells were treated with the ICG solution for 1 h, rinsed with hepatocyte-maturation medium, and then observed using a bright-field microscope (Olympus, Tokyo, Japan). After 24 h, cells were observed again to visualize excretion capability. To calculate the ICG positive area, we used ImageJ ver. 1.52m (74) software program (National Institutes of Health, Bethesda, MD, USA).

#### Image acquisition

Each sample containing cells was placed on the stage of a Nikon ECLIPSE Ti inverted fluorescence microscope equipped with a CFI plan fluor 10×/0.30 N.A. objective lens (Nikon, Tokyo, Japan), a CCD camera (ORCA-R2; Hamamatsu Photonics, Hamamatsu City, Japan), a mercury lamp (Intensilight; Nikon), an XYZ automated stage (Ti-S-ER motorized stage with encoders; Nikon), and filter cubes for fluorescence channels (DAPI and GFP HYQ; Nikon). For image acquisition, the exposure times were set at 200 ms for DAPI, 200 ms for GFP HYQ for A1AT and CYP3A7, and 800 ms for CYP3A4 and ALB.

#### Single-cell profiling based on microscopic images

Following the microscopic image acquisition, the CellProfiler software (Broad Institute of Harvard and MIT, Version 4.0.4) was used to identify cells with Otsu’s method. The fluorescence signals of individual cells were quantified automatically.

#### Periodic Acid Schiff (PAS) staining

PAS staining kit (Merck, Tokyo, Japan) were used, following the manufacture’s protocol. To calculate the PAS-stained cell area, ImageJ ver. 1.52m software program was used.

#### RNA purification

RNA was purified from cells with RNeasy Mini Kit (Qiagen, Hilden, Germany). Cells were directly lysed by adding 350 μL of lysis buffer in the kit and transfer 1.5 mL tube. Subsequently, 350 μL of 70% (v/v) ethanol was added to the tubes. Each solution was transferred to an RNeasy Mini spin column placed in a 2-mL collection tube. The column was centrifuged for 15 s at 8000 × g, and the flow-through was discarded. Then, 350 μL of buffer RW1 was added to the columns and centrifuged. Then, 80 μL of DNase digestion buffer was added to the column and incubated at 25°C for 15 min. Next, 350 μL of buffer RW1 was added to the column tube and centrifuged again. The column was washed with 500 μL of buffer RPE three times, placed in a new 2-mL tube, and centrifuged. The column was placed in a new 1.5-mL collection tube, and 50 μL of RNase-free water was added to the column; this was followed by centrifugation for 1 min at 8000× g to elute RNA into the collection tube. The RNA quality was evaluated with Agilent 2100 Bioanalyser (Agilent Technologies, Inc., USA).

#### Quantitative PCR

The primer sets used in this study are shown in **Table S4**. 1 μg of total RNA from each sample was used for cDNA synthesis using the PrimeScript RT Master Mix (TaKaRa). Each PCR was carried out in a final volume of 25 μl containing 12.5 μl of 2× TB Green Premix Ex Taq II (Tli RNaseH Plus), 2 μl (0.8 μM) of forward and reverse primers, 0.5 μl of ROX Reference Dye or Dye, 7.6 μl of nuclease-free water, and 0.4 μl (50 ng/μl) of cDNA template. The qPCR cycling conditions were 95°C for 2 min, 40 cycles at 95°C for 15 s, and 60°C for 1 min. The relative mRNA expression of some genes was calculated using the *ΔΔ*Ct method, and housekeeping genes were used to normalize the transcript levels.

#### RNA amplification and sequencing

Precisely 40 ng of total RNA was diluted with 9 μL of RNase free water, then mixed with VN primer (Oxford NANOPORE Technologies, UK) and 1 μL of 10 mM dNTPs (New England Biolabs Inc. Ipswich, Massachusetts, USA), and incubated at 65°C for 5 min to prepare the cDNA library. Separately, 4 μL of 5x RT Buffer (Thermo Fisher Scientific), 1 μL of RNaseOUT (Thermo Fisher Scientific), 1 μL of Nuclease-free water, and 2 μL of Strand-Switching Primer (Oxford NANOPORE Technologies) was mixed as the strand-switching buffer. The two solutions were mixed at 42°C for 2 min; then, 1 μL of Maxima H Minus Reverse Transcriptase (Thermo Fisher Scientific) was added. The mixture was incubated at 42°C for 90 min, 85°C for 5 min, and stored at 4°C until use as the cDNA library. Exactly 5 μL aliquot of the cDNA library solution was mixed with 25 μL of 2x LongAmp Taq Master Mix (New England Biolabs Inc.), 1.5 μL of Barcode Primers (Oxford NANOPORE Technologies), and 18.5 μL of nuclease-free water. PCR was performed (95°C for 30 sec, 18 cycles of 95°C for 15 sec, 62°C for 15 sec and 65°C for 50 sec, and then 65°C for 6 min) to barcode the cDNA for multiplexing. PCR products were stored at 4°C until use. Then, 1 μL of NEB Exonuclease 1 (New England Biolabs Inc.) was added before incubation at 37°C for 15 min, followed by incubation at 80°C for 15 min. Using Agencourt AMPure XP beads (BECKMAN COULTER Life Sciences, Indianapolis, IN), amplified DNA was purified and collected in 12 μL of Elution Buffer (Oxford NANOPORE Technologies). BioAnalyzer 2100 with High Sensitivity DNA Kit (Agilent Technologies) was used to evaluate barcoded cDNA’s amount and quality. Then, 50 fmol of the barcoded cDNA was incubated with 1 μl of Rapid Adapter to make up a total volume of 11 μL, which was incubated for 5 min at 25°C. For Nanopore sequencing, 12 μL of the prepared DNA library was mixed with 37.5 μL of Sequencing Buffer (Oxford NANOPORE Technologies) and 25.5 μL of Loading Buffer (Oxford NANOPORE Technologies). This solution was applied to the Nanopore Flow Cell (v9.4.1) and run for 24 h.

#### mRNA-seq analysis

Sequenced reads were processed and demultiplexed using the EPI2ME software (Oxford NANOPORE Technologies). Failed reads were discarded, and Fast5 files were converted into FASTQ and FASTA files using the EPI2ME software. Generated FASTQ files were loaded to BioJupies (75) for alignment with human genomes and annotation. Expressed genes were counted using BioJupies (**Table S3**), and the counts were analyzed using TCC (76) in R Bioconductor. Briefly, the counted dataset was introduced into the TCC-GUI package. Gene counts were normalized using TMM (Trimmed mean of M values) methods (77) in the edgeR package (78) with the following parameters: Filtering Threshold for Low Count Genes 30, Number of Iterations = 3; FDR < 0.1; Elimination of Potential DEGs = 0.05. Time-course RNA sequence analysis was conducted according to the R package microarray Significant Profiles (maSigPro) (version1.60.0) (79) using normalized expression value. GO analysis for DEGs was performed with the WEB-based Gene Set Analysis Toolkit (WebGestalt (24)). ‘Biological Process noRedundant,’ ‘Cellular component noRedundant’ and ‘molecular function noRedundant’ were selected for the database, and ‘genome protein-coding’ genes were selected the reference set. Also, ‘KEGG’ and ‘Reactome’ were selected for the database. The protein-coding genes among the DEGs were used as input.

#### Data availability (RNA-seq)

The mRNA-seq data have been deposited in the NCBI Gene Expression Omnibus under accession number GSE172227.

#### Statistical analysis

All experiments were carried out at least four times independently. Statistical analysis was performed with Microsoft excel. The results for the CYP3A4 assay and albumin secretion assay are presented as means ± SD. In others, where applicable, center lines of box plots indicate medians; box limits indicate the 25th and 75th percentiles as determined by R software; whiskers extend 1.5 times the interquartile range from the 25th and 75th percentiles. Comparisons between the two groups were conducted using a two-tailed Student’s *t*-test. Statistical significance was accepted for values of p < 0.05 in the present study. Comparison of CYP3A7, CYP3A4, ALB, and A1AT positive cells between two groups was conducted using the Two-sample Kolmogorov-Smirnov test followed by the Wilcoxon rank-sum test.

## Supporting information

Supplemental informations

## Acknowledgments

Funding was generously provided by the Japan Society for the Promotion of Science (JSPS; 16K14660, 17H02083, 18KK0306 and 19H02572), the Japan Agency for Medical Research and Development (AMED; 17937667) and the LiaoNing Revitalization Talents Program (XLYC1902061). The WPI-iCeMS is supported by the World Premier International Research Center Initiative (WPI), MEXT, Japan.

## Notes

**Competing Interest Statement:** The authors declare no competing interest.

### Competing Interest Statement

The authors have declared no competing interest.

## References

1. V. Y. Soldatow, E. L. Lecluyse, L. G. Griffith, I. Rusyn, In vitro models for liver toxicity testing. Toxicol. Res. (Camb). 2, 23–39 (2013).

2. J. Saito, et al., High content analysis assay for prediction of human hepatotoxicity in HepaRG and HepG2 cells. Toxicol. Vitr. 33, 63–70 (2016).

3. W. M. A. Westerink, W. G. E. J. Schoonen, Cytochrome P450 enzyme levels in HepG2 cells and cryopreserved primary human hepatocytes and their induction in HepG2 cells. Toxicol. Vitr. 21, 1581–1591 (2007).

4. C. W. Scott, M. F. Peters, Y. P. Dragan, Human induced pluripotent stem cells and their use in drug discovery for toxicity testing. Toxicol. Lett. 219, 49–58 (2013).

5. J. A. Thomson, Embryonic Stem Cell Lines Derived from Human Blastocysts. Science (80-.). 282, 1145–1147 (1998).

6. K. Takahashi, et al., Induction of Pluripotent Stem Cells from Adult Human Fibroblasts by Defined Factors. Cell 131, 861–872 (2007).

7. L. T. Ang, et al., A Roadmap for Human Liver Differentiation from Pluripotent Stem Cells. Cell Rep. 22, 2190–2205 (2018).

8. Y. Avior, et al., Microbial-Derived Lithocholic Acid and Vitamin K2 Drive the Metabolic Maturation of Pluripotent Stem Cells-Derived and Fetal Hepatocytes. Hepatology 62, 265–278 (2015).

9. A. Carpentier, et al., Hepatic differentiation of human pluripotent stem cells in miniaturized format suitable for high-throughput screen. Stem Cell Res. 16, 640–650 (2016).

10. D. Zhao, et al., Promotion of the efficient metabolic maturation of human pluripotent stem cell-derived hepatocytes by correcting specification defects. Cell Res. 23, 157–161 (2013).

11. K. Si-Tayeb, et al., Highly efficient generation of human hepatocyte-like cells from induced pluripotent stem cells. Hepatology 51, 297–305 (2010).

12. P. Baconnier, G. Benchetrit, M. Tanche, Liver heat production and temperature regulation in the anesthetized dog. Am. J. Physiol. Integr. Comp. Physiol. 237, R334–R339 (1979).

13. J. Simcox, et al., Global Analysis of Plasma Lipids Identifies Liver-Derived Acylcarnitines as a Fuel Source for Brown Fat Thermogenesis. Cell Metab. 26, 509–522.e6 (2017).

14. M. E. Hossain, et al., Direct exposure to mild heat promotes proliferation and neuronal differentiation of neural stem/progenitor cells in vitro. PLoS One 12, 1–17 (2017).

15. T. Yamaguchi, T. Suzuki, H. Arai, S. Tanabe, Y. Atomi, Continuous mild heat stress induces differentiation of mammalian myoblasts, shifting fiber type from fast to slow. Am. J. Physiol. - Cell Physiol. 298, 140–148 (2010).

16. M. Nakamura, et al., Temperature sensitivity of human spermatogonia and spermatocytes in vitro. Syst. Biol. Reprod. Med. 19, 127–132 (1987).

17. K. Kamei, M. Yoshioka, S. Terada, Y. Tokunaga, Y. Chen, Three-dimensional cultured Liver-on-a-Chip with mature hepatocyte-like cells derived from human pluripotent stem cells.

18. C. M. Ho, et al., Use of indocyanine green for functional assessment of human hepatocytes for transplantation. Asian J. Surg. (2012) https:/doi.org/10.1016/j.asjsur.2012.04.017.

19. W. De Graaf, et al., Transporters involved in the hepatic uptake of 99mTc-mebrofenin and indocyanine green. J. Hepatol. 54, 738–745 (2011).

20. M. Kusano, N. Kokudo, M. Toi, M. Kaibori, ICG Fluorescence Imaging and Navigation Surgery, M. Kusano, N. Kokudo, M. Toi, M. Kaibori, Eds. (Springer Japan, 2016) https:/doi.org/10.1007/978-4-431-55528-5.

21. R. Pal, M. K. Mamidi, A. K. Das, R. Bhonde, Diverse effects of dimethyl sulfoxide (DMSO) on the differentiation potential of human embryonic stem cells. Arch. Toxicol. 86, 651–661 (2012).

22. G. Stelzer, et al., The GeneCards suite: From gene data mining to disease genome sequence analyses. Curr. Protoc. Bioinforma. 2016, 1.30.1–1.30.33 (2016).

23. T. Matsunaga, et al., Mechanisms of CYP3A induction by glucocorticoids in human fetal liver cells. Drug Metab. Pharmacokinet. 27, 653–657 (2012).

24. J. Wang, S. Vasaikar, Z. Shi, M. Greer, B. Zhang, WebGestalt 2017: A more comprehensive, powerful, flexible and interactive gene set enrichment analysis toolkit. Nucleic Acids Res. 45, W130–W137 (2017).

25. L. Fagerberg, et al., Analysis of the Human Tissue-specific Expression by Genome-wide Integration of Transcriptomics and Antibody-based Proteomics. Mol. Cell. Proteomics 13, 397–406 (2014).

26. M. Inada, et al., Stage-specific regulation of adhesion molecule expression segregates epithelial stem/progenitor cells in fetal and adult human livers. Hepatol. Int. 2, 50–62 (2008).

27. M. J. Williams, A. D. Clouston, S. J. Forbes, Links between hepatic fibrosis, ductular reaction, and progenitor cell expansion. Gastroenterology 146, 349–356 (2014).

28. D. Schuppan, M. Ashfaq-Khan, A. T. Yang, Y. O. Kim, Liver fibrosis: Direct antifibrotic agents and targeted therapies. Matrix Biol. 68-69, 435–451 (2018).

29. M. A. Karsdal, et al., The good and the bad collagens of fibrosis – Their role in signaling and organ function. Adv. Drug Deliv. Rev. 121, 43–56 (2017).

30. N. Carolina, E. Thomas, J. Brown, V. Affairs, Functional Maturation of Induced Pluripotent Stem Cell Hepatocytes in Extracellular Matrix—A Comparative Analysis of Bioartificial Liver Microenvironments. Stem Cells Transl. Med. 5, 1257–1267 (2016).

31. S. Nakai, et al., Collagen vitrigel promotes hepatocytic differentiation of induced pluripotent stem cells into functional hepatocyte-like cells. Biol. Open 8, bio042192 (2019).

32. B. Hildebrandt, The cellular and molecular basis of hyperthermia. Crit. Rev. Oncol. Hematol. 43, 33–56 (2002).

33. F. Soetaert, P. Korangath, D. Serantes, S. Fiering, R. Ivkov, Cancer therapy with iron oxide nanoparticles: Agents of thermal and immune therapies. Adv. Drug Deliv. Rev. 163-164, 65–83 (2020).

34. K. B. Laupland, Fever in the critically ill medical patient. Crit. Care Med. 37, S273–S278 (2009).

35. J. A. Majda, E. W. Gerner, B. Vanlandingham, K. R. Gehlsen, A. E. Cress, Heat shock-induced shedding of cell surface integrins in A549 human lung tumor cells in culture. Exp. Cell Res. 210, 46–51 (1994).

36. Z. T. Gu, et al., Heat stress induces apoptosis through transcription-independent p53-mediated mitochondrial pathways in human umbilical vein endothelial cell. Sci. Rep. 4, 1–10 (2014).

37. G. Stelzer, et al., The GeneCards Suite: From Gene Data Mining to Disease Genome Sequence Analyses. Curr. Protoc. Bioinforma. 54, 1.30.1–1.30.33 (2016).

38. C.-X. Yuan, W. B. Gurley, Potential targets for HSF1 within the preinitiation complex. Cell Stress Chaperones 5, 229 (2000).

39. D.-P. Li, S. Periyasamy, T. J. Jones, E. R. Sánchez, Heat and Chemical Shock Potentiation of Glucocorticoid Receptor Transactivation Requires Heat Shock Factor (HSF) Activity. J. Biol. Chem. 275, 26058–26065 (2000).

40. T. J. Jones, et al., Enhancement of Glucocorticoid Receptor-Mediated Gene Expression by Constitutively Active Heat Shock Factor 1. Mol. Endocrinol. 18, 509–520 (2004).

41. C. Bonnans, J. Chou, Z. Werb, Remodelling the extracellular matrix in development and disease. Nat. Rev. Mol. Cell Biol. 15, 786–801 (2014).

42. O. Levi-Galibov, et al., Heat Shock Factor 1-dependent extracellular matrix remodeling mediates the transition from chronic intestinal inflammation to colon cancer. Nat. Commun. 11, 6245 (2020).

43. A. M. Reimold, et al., An essential role in liver development for transcription factor XBP-1. Genes Dev. 14, 152–157 (2000).

44. K. Zhang, et al., The unfolded protein response sensor IRE1α is required at 2 distinct steps in B cell lymphopoiesis. J. Clin. Invest. 115, 268–281 (2005).

45. S. Rachidi, et al., Endoplasmic reticulum heat shock protein gp96 maintains liver homeostasis and promotes hepatocellular carcinogenesis. J. Hepatol. 62, 879–888 (2015).

46. G. Aran, et al., CD5L is upregulated in hepatocellular carcinoma and promotes liver cancer cell proliferation and antiapoptotic responses by binding to HSPA5 (GRP78). FASEB J. 32, 3878–3891 (2018).

47. S. H. Choi, et al., MMP9 processing of HSPB1 regulates tumor progression. PLoS One 9(2014).

48. M. Uhlén, et al., Tissue-based map of the human proteome. Science (80-.). 347(2015).

49. P. J. Thul, et al., A subcellular map of the human proteome. Science (80-.). 356(2017).

50. M. Uhlen, et al., A pathology atlas of the human cancer transcriptome. Science (80-.). 357(2017).

51. R. D. Ye, L. Sun, Emerging functions of serum amyloid A in inflammation. J. Leukoc. Biol. 98, 923–929 (2015).

52. Y. Wang, et al., Serum amyloid a induces M2b-like macrophage polarization during liver inflammation. Oncotarget 8, 109238–109246 (2017).

53. A. Das, et al., Monocyte and Macrophage Plasticity in Tissue Repair and Regeneration. Am. J. Pathol. 185, 2596–2606 (2015).

54. Y. Watanabe, et al., Mesenchymal Stem Cells and Induced Bone Marrow-Derived Macrophages Synergistically Improve Liver Fibrosis in Mice. Stem Cells Transl. Med. 8, 271–284 (2019).

55. T. Touboul, et al., Generation of functional hepatocytes from human embryonic stem cells under chemically defined conditions that recapitulate liver development. Hepatology 51, 1754–1765 (2010).

56. H. C. Bisgaard, P. P. T. Nagy Ton, Z. Hu, S. S. Thorgeirsson, Modulation of keratin 14 and ?-fetoprotein expression during hepatic oval cell proliferation and liver regeneration. J. Cell. Physiol. 159, 475–484 (1994).

57. E. Schmelzer, et al., Human hepatic stem cells from fetal and postnatal donors. J. Exp. Med. 204, 1973–1987 (2007).

58. Y.-R. Lou, A. W. Leung, Next generation organoids for biomedical research and applications. Biotechnol. Adv. 36, 132–149 (2018).

59. T. Takebe, et al., Vascularized and functional human liver from an iPSC-derived organ bud transplant. Nature 499, 481–484 (2013).

60. S. Kiyonaka, et al., Genetically encoded fluorescent thermosensors visualize subcellular thermoregulation in living cells. Nat. Methods 10, 1232–1238 (2013).

61. M. Nakano, et al., Genetically encoded ratiometric fluorescent thermometer with wide range and rapid response. PLoS One 12, e0172344 (2017).

62. K. Okabe, R. Sakaguchi, B. Shi, S. Kiyonaka, Intracellular thermometry with fluorescent sensors for thermal biology. Pflügers Arch. - Eur. J. Physiol. 470, 717–731 (2018).

63. H. Kimura, Y. Sakai, T. Fujii, Organ/body-on-a-chip based on microfluidic technology for drug discovery. Drug Metab. Pharmacokinet. 33, 43–48 (2018).

64. K. H. Benam, et al., Engineered In Vitro Disease Models. Annu. Rev. Pathol. Mech. Dis. 10, 195–262 (2015).

65. J. H. Sung, et al., Recent Advances in Body-on-a-Chip Systems. Anal. Chem. 91, 330–351 (2019).

66. A. Soto-Gutierrez, A. Gough, L. A. Vernetti, D. L. Taylor, S. P. Monga, Pre-clinical and clinical investigations of metabolic zonation in liver diseases: The potential of microphysiology systems. Exp. Biol. Med. 242, 1605–1616 (2017).

67. K. Kamei, et al., Integrated heart/cancer on a chip to reproduce the side effects of anti-cancer drugs in vitro. RSC Adv. 7, 36777–36786 (2017).

68. K. Kamei, M. Yoshioka, S. Terada, Y. Tokunaga, Y. Chen, Three-dimensional cultured liver-on-a-Chip with mature hepatocyte-like cells derived from human pluripotent stem cells. Biomed. Microdevices 21, 73 (2019).

69. L.-D. Ma, et al., Design and fabrication of a liver-on-a-chip platform for convenient, highly efficient, and safe in situ perfusion culture of 3D hepatic spheroids. Lab Chip 18, 2547–2562 (2018).

70. E. Moradi, S. Jalili-Firoozinezhad, M. Solati-Hashjin, Microfluidic organ-on-a-chip models of human liver tissue. Acta Biomater. 116, 67–83 (2020).

71. N. Tsamandouras, et al., Integrated Gut and Liver Microphysiological Systems for Quantitative In Vitro Pharmacokinetic Studies. AAPS J. 19(2017).

72. C. H. Amon, et al., MEMS-enabled thermal management of high-heat-flux devices EDIFICE: Embedded droplet impingement for integrated cooling of electronics. Exp. Therm. Fluid Sci. 25, 231–242 (2001).

73. Y. Cui, et al., MEMS-based dual temperature control measurement method for thermoelectric properties of individual nanowires. MRS Commun. 10, 620–627 (2020).

74. C. A. Schneider, W. S. Rasband, K. W. Eliceiri, NIH Image to ImageJ: 25 years of image analysis. Nat. Methods 9, 671–675 (2012).

75. A. M. Denis Torre, Alexander Lachmann, BioJupies: Automated Generation of Interactive Notebooks for RNA-seq Data Analysis in the Cloud. Physiol. Behav. 176, 139–148 (2019).

76. W. Su, J. Sun, K. Shimizu, K. Kadota, TCC-GUI: A Shiny-based application for differential expression analysis of RNA-Seq count data. BMC Res. Notes 12, 1–6 (2019).

77. M. D. Robinson, A. Oshlack, A scaling normalization method for differential expression analysis of RNA-seq data. Genome Biol. 11(2010).

78. M. D. Robinson, D. J. McCarthy, G. K. Smyth, edgeR: A Bioconductor package for differential expression analysis of digital gene expression data. Bioinformatics 26, 139–140 (2009).

79. A. Conesa, M. J. Nueda, A. Ferrer, M. Talón, maSigPro: A method to identify significantly differential expression profiles in time-course microarray experiments. Bioinformatics 22, 1096–1102 (2006).

