## Supplemental informations for "Heat treatment functionalizes hepatocyte-like cells derived from human embryonic stem cells"

Address : Yoshida-Ushinomiya-cho, Sakyo-ku, Kyoto, 606-8501, JAPAN

**This PDF file includes:**

Figures S1 to S4

Tables S1 to S4

**Fig. S1**

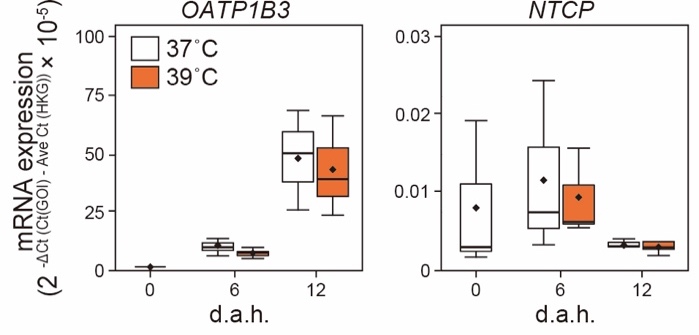

Gene expression of solute carrier organic anion transporter family member 1B3 (*OATP1B3*) and Na^+^-taurocholate cotransporting polypeptide (*NCTP*), in 37˚C- and 39˚C-treated hESC-HLCs at 0, 6 and 12 d.a.h. evaluated by quantitative RT-PCR (n = 4).

**Fig. S2**

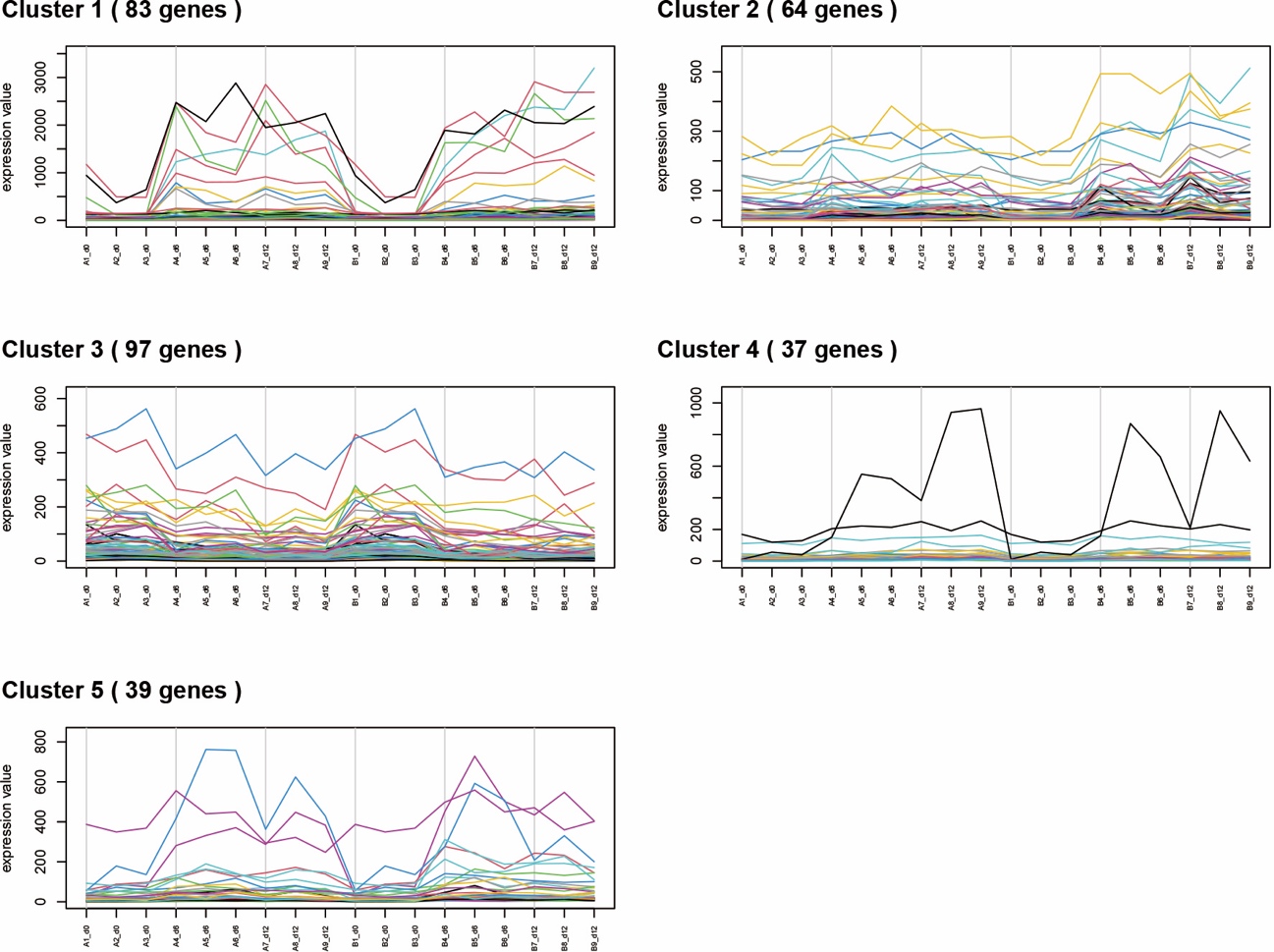

Five clusters of the specific gene signatures in hESC-HLCs with heat treatments at 37˚C and 39˚C by global transcriptional analysis to identify.

**Fig. S3**

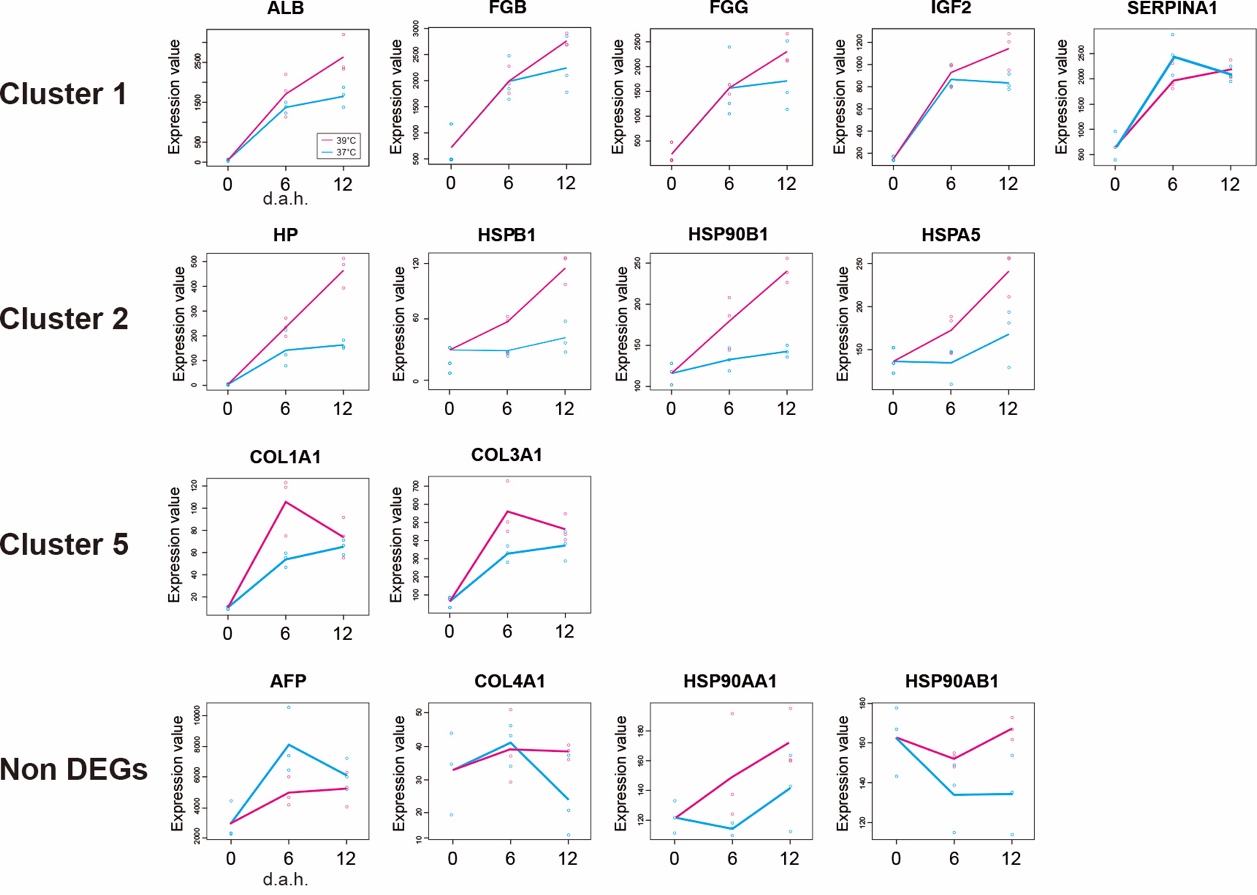

Time-course plots of typical genes in Cluster 1, 2, 5, and non DEGs.

**Fig. S4**

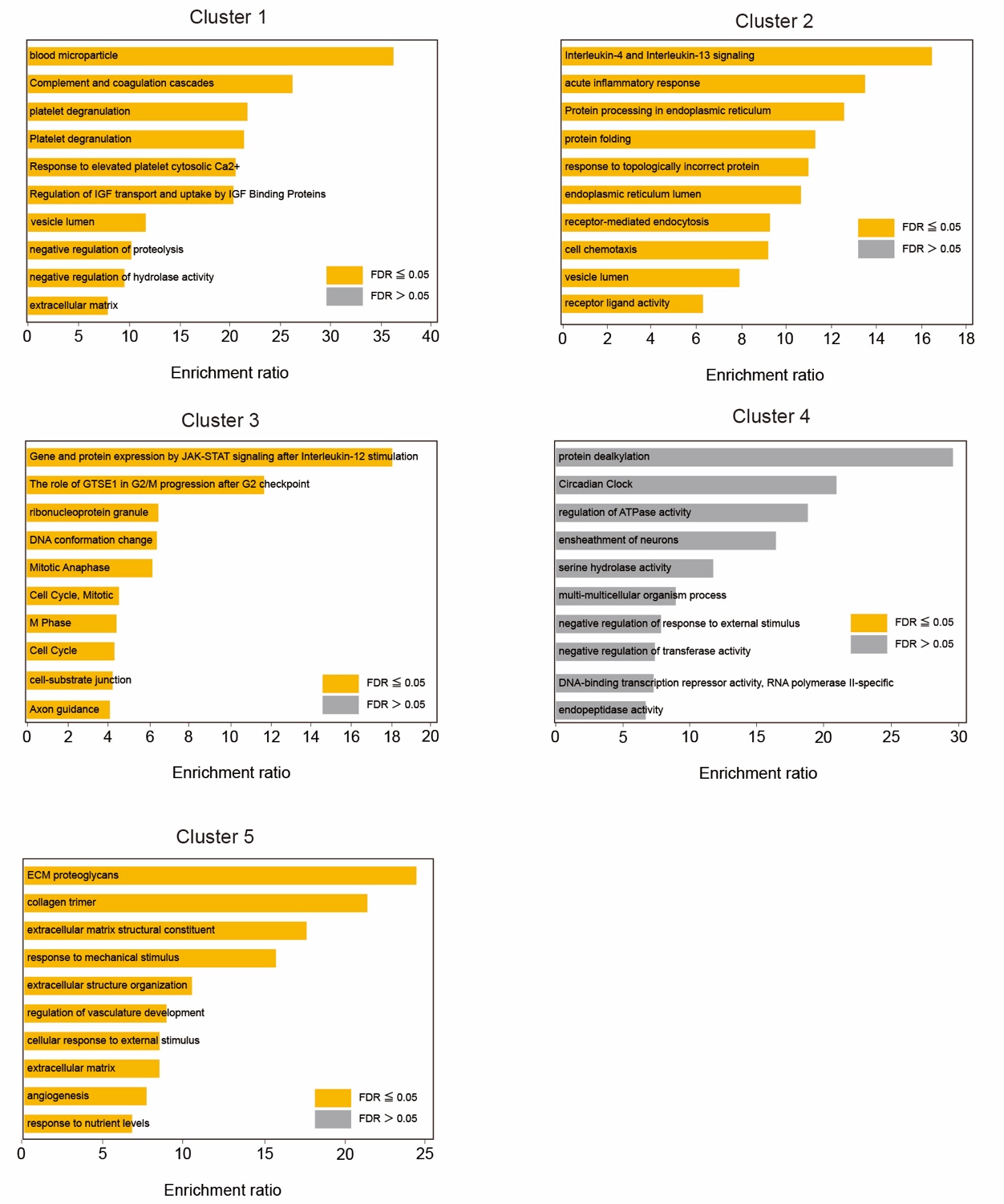

Enriched GO, KEGG and Reactome terms of differentially expressed genes in 39˚C-treated hESC-HLCs.

**Table S1.** Gene counts by RNA-seq for 37˚C- and 39˚C-treated hESC-HLCs at 0, 6, and 12 days after heat treatment (d.a.h.).

**Table S2.** List of differentially expressed genes (DEGs) between 37˚C- and 39˚C-treated hESC-HLCs.

**Table S3.** List of gene ontological terms obtained with DEGs between 37˚C- and 39˚C-treated hESC-HLCs.

**Table S4.** List of primers for quantitative RT-PCR in this study.

**S.I. References**

Use the "Insert Citation" button to add citations to this document.
